# The effects of long-term atorvastatin use on the cytoarchitecture of new and older neurons of sensorimotor nucleus HVC in a juvenile songbird model

**DOI:** 10.1101/196204

**Authors:** Shuk C. Tsoi, Alicia C. Barrientos, Carolyn L. Pytte, David S. Vicario, Mimi L. Phan

**Affiliations:** Biology Dept., The Graduate Center, CUNY, New York, NY; Psychology Dept., The Graduate Center, CUNY, New York, NY; Psychology Dept., Queens College, New York, NY; Psychology Dept., Rutgers-The State University of New Jersey, New Brunswick, NJ

## Introduction

Statins are a commonly-used class of cholesterol-lowering drugs which effectively reduce the production of plasma cholesterol by the liver, and thus are associated with reducing the risk of cardiovascular diseases (Horwich et al., 2004). However, cholesterol is also produced *de novo* in the brain, raising questions of whether oral statins that cross from the periphery into the brain impact brain cholesterol, neural tissue, and cognition (Eckert et al., 2001, Fong, 2014, Banach et al., 2017). The vast majority of research on statins and cognition convincingly demonstrate that many statins are neuroprotective and ameliorate cognitive decline in patients with neurodegenerative disease and brain injuries, e.g., (Bunt and Hogan, 2017, Chan et al., 2017). However, much less is known about whether and how statins impact neural function in patients who are neurally healthy and are being treated only for high levels of plasma cholesterol (Wagstaff et al., 2003, Samaras et al., 2016, Suraweera et al., 2016). The results of studies investigating effects of statins on healthy brains are inconsistent, but have led to FDA warnings on statins indicating potential cognitive impairment with use (https://www.fda.gov/Drugs/DrugSafety/ucm293101.htm). Moreover, almost no work has been conducted on neural effects of statin use in neurally healthy juveniles, either in pediatric populations or animal models (Wiegman et al., 2004, Schreurs, 2010). Because neurogenesis requires continual formation of cholesterol-rich lipid membranes, we speculated that one likely candidate for statin-related neural disruption is the process of neurogenesis and the integrity of membranes formed specifically during statin use. Thus, we investigated the effects of high doses of atorvastatin (Lipitor^®^) administered to juvenile songbirds on the cytoarchitecture of newborn neurons *in vivo*.

Post embryonic brain cholesterol is mainly produced by astrocytes and oligodendrocytes, whereas embryonic brain cholesterol is primarily produced by microglia; in both cases synthesis is regulated independently from that of blood cholesterol (Saher et al., 2005, Vance et al., 2005, Funfschilling et al., 2007, Orth and Bellosta, 2012). Neurons can also produce cholesterol, although do so much more slowly than do glia (Nieweg et al., 2009). The majority (∼70%) of brain cholesterol is found in the myelin sheath with the remainder primarily incorporated into glial and neuronal membranes (Dietschy and Turley, 2001, 2004, Goluszko and Nowicki, 2005, Dietschy, 2009).

The plasma membrane incorporates cholesterol into specialized subdomains called lipid rafts (Chaudhuri and Chattopadhyay, 2011), which play a critical role not only in membrane structure and organization, but also membrane dynamics and interactions with transmembrane proteins (Liscum and Underwood, 1995, Simons and Ikonen, 2000, Mouritsen and Zuckermann, 2004). Altering concentrations of neuronal membrane cholesterol *in vitro* has been shown to affect acetylcholine receptor function, as well as the enzyme ATPase that stimulates conformational changes of the sodium-potassium pump (Yeagle, 1989). Intracellular cholesterol is important for proper synapse formation and axonal growth (Pfrieger, 2003a, b). Cholesterol also plays a role in the transport of proteins and paracrine signaling (Goluszko and Nowicki, 2005, Vance et al., 2005, Piomelli et al., 2007, Orth and Bellosta, 2012, Vance, 2012).

Treating neurons with statins *in vitro* has resulted in a substantial list of effects on cell structure including damaged organelle morphology, fragmented neurites (Pavlov et al., 1995), compacted nuclei with condensed chromatin (Pavlov et al., 1995, Tanaka et al., 2000), reduced synapse densities (Mailman et al., 2011), spine shrinkage, and a decrease in neurite density (Schulz et al., 2004). Statin treatment has also impaired neuronal function, decreasing synaptic vesicle release in primary culture of rat hippocampal neurons following one week of lovastatin treatment (Mailman et al., 2011). In addition, statin exposure, or otherwise blocking cholesterol production, has induced cell death of cortical neurons (Michikawa and Yanagisawa, 1999, Tanaka et al., 2000) and cerebellar neurons (Marz et al., 2007, Xiang and Reeves, 2009). In addition to effects on neurons, *in vitro* statin exposure altered cellular morphology and decreased survival of astrocytes (Pavlov et al., 1995), microglia (Marz et al., 2007), and oligodendrocytes (Xiang and Reeves, 2009). Oral statin use (lovastatin) has been shown to lower brain membrane cholesterol in normal young-adult and middle-aged adult mice (Eckert et al., 2001). However, effects of statins specifically on cell structure, function, and survival *in vivo* have not been assessed.

Oral statins have been shown act in the brain and decrease available brain cholesterol *in vivo*. However, which statins do this by crossing the blood-brain barrier (BBB), or by permeating non-BBB entry points, is not clear (Locatelli et al., 2002, Kirsch et al., 2003, Burns et al., 2006, Fong, 2014). Regardless of the mechanism of entry, regions of the brain that undergo life-long neurogenesis, requiring a high demand for new membrane production, may be particularly sensitive to alterations in brain cholesterol. Neural stem and progenitor cells are dependent on endogenous cholesterol synthesis for survival and cholesterol derivatives have been shown to directly contribute to neurogenesis and neuronal survival during embryonic brain development (Theofilopoulos and Arenas, 2015). In addition, genetically reduced activity of a cholesterol biosynthesis enzyme caused abnormal neuroprogenitor fate determination, among other alterations of embryonic neurogenesis (Driver et al., 2016). Thus, dynamics of adult neurogenesis, and in particular, membrane structure, may be distinctly vulnerable to statin use.

Currently, four statins are approved for pediatric populations suffering from early onset obesity and a genetic predisposition for developing abnormally high levels of blood plasma cholesterol (familial hypercholesterolemia, FH). Moreover, health care professionals can use statins for off-label purposes and overextend statin treatment to children who have high levels of serum cholesterol but do not have the genetic disorder (Gelissen et al., 2014). However, studies on pediatric statin users have solely focused on peripheral effects of organ toxicity, pharmacokinetics, body growth and sexual maturation, and lack neural examinations. Most cognitive research in statin treated children has focused on subjects with neurofibromatosis type 1 or other nervous system disorders. Research on effects of statins on cognitive function or development in healthy children appears to be limited to a single study of their school performance over a course of 2 years (Wiegman et al., 2004), which, although laudatory, lacks details of cognitive assessment. Likewise, studies of the effects of statins on brain structure in juvenile animal models are scarce (Wise et al., 2011). As a first step toward understanding the effects of statins on neural tissue in juveniles, we used a songbird model system and focused on determining whether statins altered new neuron numbers and cytoarchitecture.

## Materials and methods

### Animals

All work was approved by the Institutional Animal Care and Use Committees of Queens College, CUNY, and Rutgers University. Zebra finches were bred and housed in the Rutgers University vivarium. The birds were kept on a 12 hr light, 12 hr dark cycle with *ad libitum* access to food and water. As part of a study on the effects of stains on song learning and memory, only male birds were selected for this work as only male zebra finches produce song.

### Atorvastatin-treatment

Starting at day 43, experimental birds were given a daily dose of 40 mg/kg of atorvastatin (Lipitor^®^, Pfizer) dissolved in 50 μl of water, administered by pipette (n=6). Controls were given the same volume of water vehicle (n=7). Treatments continued daily until perfusion at 110 days of age (67 days of treatment).

### BrdU administration

New neurons were quantified in the nucleus HVC region of the song system that functions in song learning and production approximately one month post cell birthdating. To label mitotically active cells, birds received intramuscular injections of the thymidine analog 5-bromo-2’-deoxyuridine (BrdU) three times a day for three consecutive days (78 microliters of a 10 mg/ml solution in 0.1 M Tris buffered saline, approximately 0.074 mg/g, Boehringer Mannheim) 30-33 days before perfusion.

### Perfusion and histology

The birds were deeply anesthetized with Euthasol (Henry Schien), then perfused with 0.1 M phosphate buffered saline (PBS) followed by 4% paraformaldehyde (PFA). The brains were removed and fixed with PFA for one hour then stored in PBS overnight at 4°C. Brains were divided into hemispheres, then dehydrated and embedded in polyethylene glycol (PEG) as in (Pytte et al., 2012). Hemispheres were sagittally sectioned at 6 μm using a rotary microtome. Every eighth section was collected in three series and mounted onto Superfrost+ charged slides and stored at −20°C.

### Immunohistochemistry

One slide series was used to label BrdU and the neuron-specific protein Hu. Slides were heated in citrate buffer for 10 min at 90-95°C then rinsed for 5 min in PBS. Then slides were then soaked in a solution of 0.1 N HCl with 0.28% pepsin at 37°C for 3 min, and rinsed three times for 5 min each in PBS. To block non-specific binding sites, slides were incubated for one hour in 0.3% Triton-X and 10% normal donkey serum in PBS. Tissue was then incubated overnight at 4°C in anti-BrdU primary antibody made in sheep (1:194 in blocking solution, Capralogics). After 3 PBS washes of 10 min each, 4 drops of avidin block (Invitrogen) were applied to each slide. Slides underwent three 10 min PBS rinses followed by 4 drops of biotin block (Invitrogen). Tissue was then incubated for 1 hour at room temperature in biotinylated donkey anti-sheep (1:200 in block, Abcam), rinsed 3 times for 10 minutes each in PBS, followed by a 2 hr incubation in streptavidin-conjugated Alexa 488 donkey anti-sheep IgG (1:800 in PBS, Invitrogen). Next, the slides were rinsed 3 times for 10 min with PBS, then incubated for 1 hr in 10% block. A primary antibody to Hu made in mouse (1:200 in PBS, Life Technologies) was applied to slides and incubated overnight. Slides were rinsed in PBS 3 times for 10 min, then treated with donkey anti-mouse conjugated to Cy3 (1:80 in block) for 1 hr. Slides were soaked in 3 PBS washes for 10 min, then submerged in deionized water for 30 sec. Finally, slides underwent dehydration in 50% ethyl alcohol (EtOH) for 30 sec, then 70% EtOH for 30 sec followed by 95% EtOH, 100% EtOH, and xylenes each for 1 min. After removal from xylenes, slides were coverslipped with krystalon.

### Microscopy

All microscopy was performed blind to bird identity and treatment. Brain tissue was observed using a computer-controlled fluorescence microscope (Olympus BX51). Mapping software (Lucivid microprojection and Neurolucida, Microbrightfield Bioscience Inc.) was used to trace HVC, mark and quantify new neurons, trace neuronal somas, and produce soma feature measurements on closed contours (described below). HVC was identified by the location of the hippocampus, cerebellum and RA under dark field. Cells in HVC were counted in 10-12 sections from ∼1600 to 2200 µm lateral to the midline of each hemisphere.

### Cell quantification

BrdU+/Hu+ cells were visualized by switching between the FITC and rhodamine filter. We quantified the density of these neurons by positioning a marker on cells expressing a BrdU+ nucleus which co-localized with a Hu+ soma. We calculated the density of BrdU+/Hu+ cells per mm^2^ in 10-12 sections of HVC. To determine whether statin treatments affected the rest of the HVC neuronal population, we also traced the contours of neurons that did not contain BrdU. This population includes neurons that were born when BrdU was not available, both during embryogenesis and in adulthood. Presumably, most of these cells were older than 30-days old. Most new neurons do not survive in the first weeks of birth in songbirds (Alvarez-Buylla and Nottebohm, 1988). Those that do, survive for many years (Walton et al., 2012). Therefore, we reasoned that the majority of non BrdU-labeled neurons that we quantified were likely older than the 30 day old BrdU-labeled cohort. We sampled HVC to quantify BrdU-/Hu+ neurons by placing a grid of 100 µm by 100 µm superimposed over each HVC tracing, and only BrdU-/Hu+ neurons from every fourth grid were counted and traced. To avoid counting errors, neurons whose somas were less than halfway in the top, left-hand corner of the sampling grid were not traced (Tsoi et al., 2014).

### Cell feature measurements

All BrdU+/HU+ cell somas, and the sample of counted BrdU-/Hu+ cell somas, were traced and feature measurements were automatically generated by Neurolucida, defined by the Neurolucida User’s Guide (linked to http://www.mbfbioscience.com/) as follows.

**Aspect Ratio** is a measure of flatness. Values closer to 1.0 indicate contours resembling a circle, and values less than 1.0 indicate increasing oval or flatness.

**Compactness** is a measure of the relationship between an outline’s area and maximum diameter, with values closer to 1.0 indicating a circle (which has a large area to perimeter relationship), whereas values less than 1.0 describe the inverse.

The **Roundness** parameter is the square of an outline’s compactness value, and it employs the same scale of 0.0 to 1.0. Squaring these values helps to differentiate cells or outlines with smaller compactness values.

**Shape Factor and Form Factor** measure the complexity of a tracing’s outline. For shape factor, values closer to 3.54 indicate a smooth contour, whereas values less than 3.54 describe an uneven surface. Form factor is a similar parameter to shape factor, except the numerical values of the properties of smoothness and roughness are measured using a different scale: values closer to 1.0 indicate unevenness, and values less than 1.0 indicate smoothness.

### Statistics

We used two-tailed t-tests to determine whether there were differences between the statin-treated birds and the control birds in the following: Numbers of new BrdU-labeled neurons in HVC (30-33 days old), numbers of new BrdU+/Hu+ neurons in HVC (∼1-3 weeks old), and neuronal soma feature measurements, using the Bonferroni correction for multiple comparisons. Values reported are means ± SEMs.

## Results

### BrdU+/Hu+ cells

Atorvastatin treatments did not affect the numbers of 30-33 day old new neurons in HVC (controls=18.943, SEM=2.614, statin-treated=18.476, SEM=2.754, t (11) = 0.12, p= 0.906, Figures 1, 2). There was also no difference between groups in soma area (controls: 57.012μ^2^, SEM=5.209; statin-treated=53.177μ^2^, SEM=2.927, t (11)=-0.584, p=0.57), or in measures of minimum or maximum diameters (p>0.05 for all). We did, however, find a difference between groups in several parameters of soma shape and surface smoothness.

**Figure 1.**
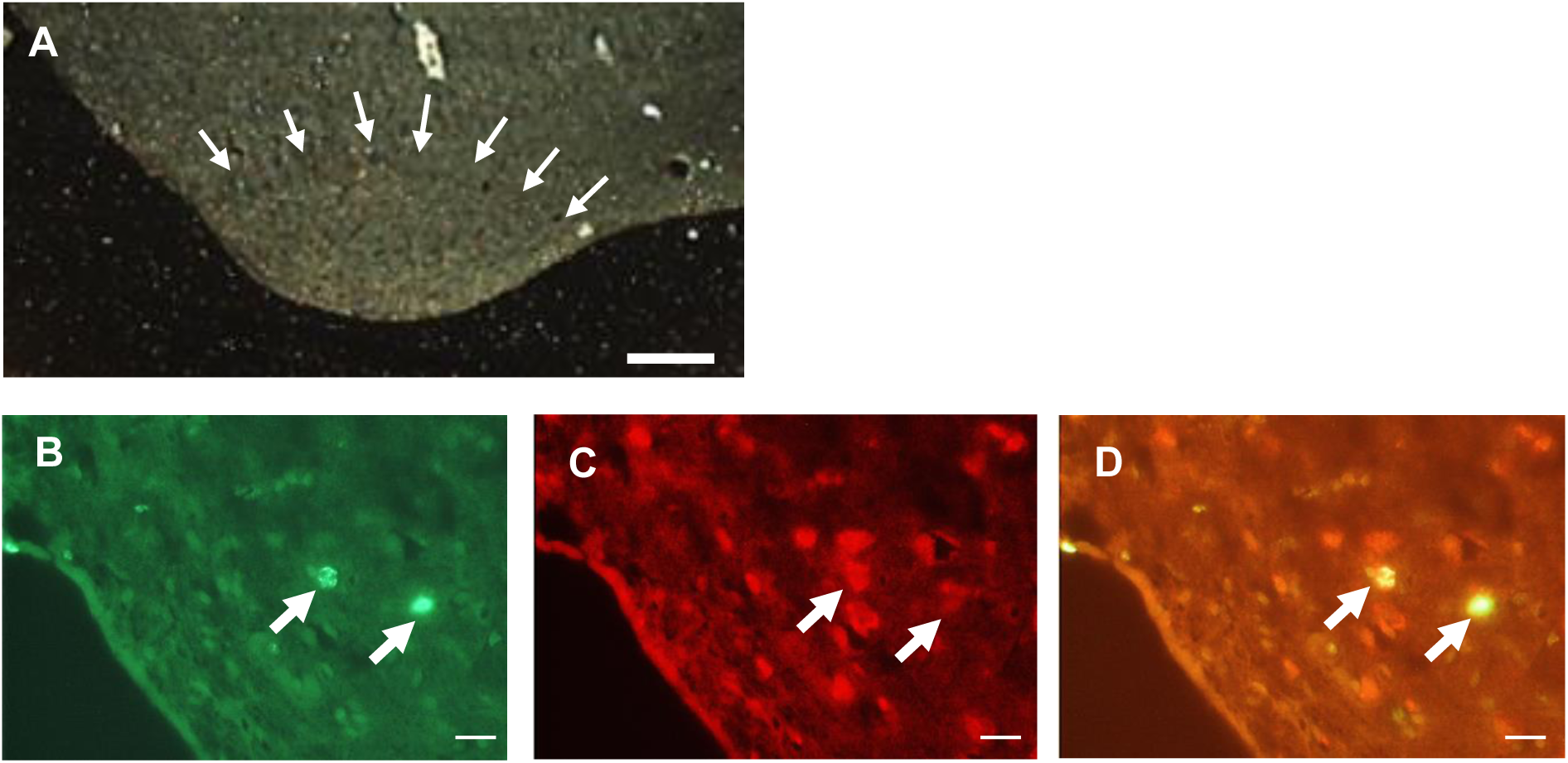
The boundary of nucleus HVC is marked by arrows, shown in dark field (A). BrdU-labeled cells visualized with a fluorescein isothiocyanate (FITC) filter (B) and Hu-labeled cells visualized with a rhodamine filter (C). Double-labeled cells were identified by alternating between these two filters and also using a dual FITC-rhodamine filter (D). Scale bars = 100 μm in A, 10 μm in B-D.

**Figure 2.**
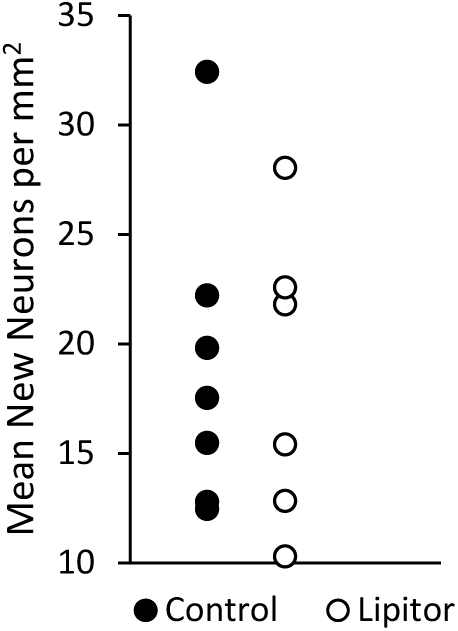
There was no difference in the density of ∼30 day old neurons between atorvastatin-treated birds and controls.

There was a difference between groups in soma aspect ratio, a measure of flatness, Figure 3). Neurons of control birds were closer to 1.0, indicating a rounder soma (0.697±0.011) whereas the neurons of atorvastatin-treated birds were lower than 1.0, indicating flatter somas (0.654±0.013). Neuronal somas of atorvastatin-treated birds were significantly flatter than controls (t (12)=-2.44, p=0.031).

**Figure 3.**
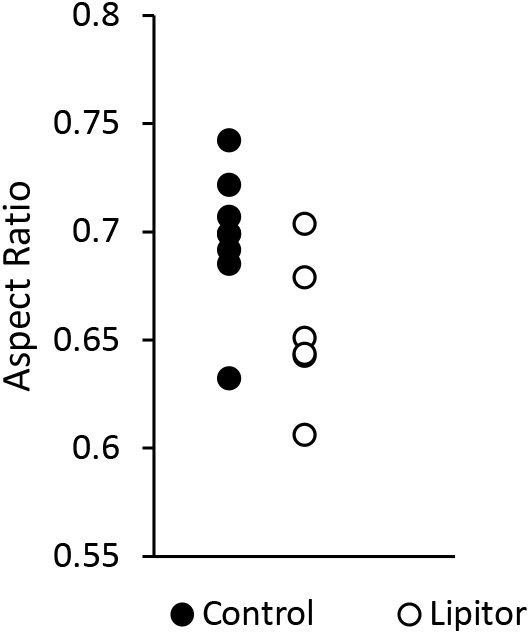
In atorvastatin treated birds, the somas of ∼30 day old neurons were flatter than in controls. Each point is the mean of cell contours for an individual bird.

There was a trend toward a difference between groups in compactness (Figure 4A), such that neurons of atorvastatin-treated birds (0.491±0.019) showed a trend towards lower compactness values relative to controls (0.536±0.014, t (12)=-1.84, p=0.086). Atorvastatin-treated birds also had neurons with lower roundness values (Figure 4B, 0.439±0.017) than controls (0.494±0.015, t (12)= 2.37, p=0.035).

**Figure 4.**
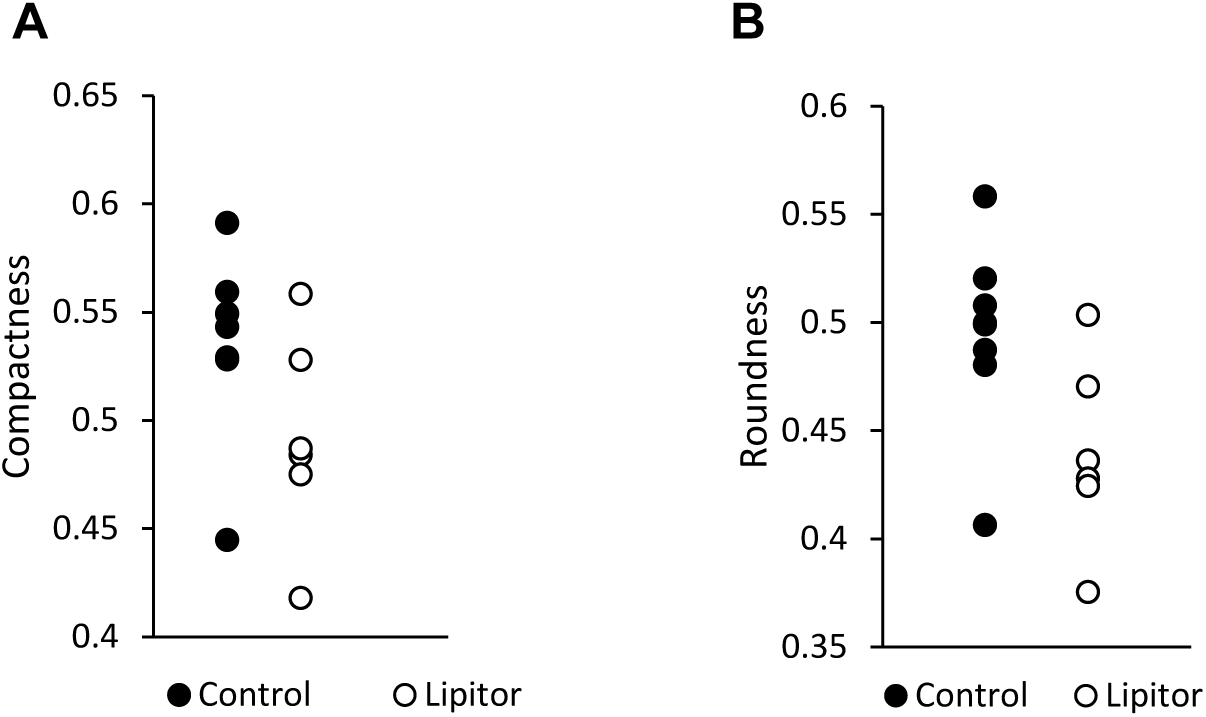
Compactness and roundness are two parameters that are closely related; the latter being the square of compactness which helps to elucidate objects with smaller compactness values. A value closer to 1.0 describes a circle. A trend was observed for compactness (A), but by squaring these values, it was found that the somas of atorvastatin treated birds have smaller roundness values relative to controls (B).

There was a difference between groups in shape factor (Figure 5A) and form factor (Figure 5B), measures of outline complexity. Neurons of control birds had higher values for shape factor (4.248±0.050) and lower values for form factor (0.717±0.014), whereas neurons of atorvastatin-treated birds had lower values for shape factor (4.06±0.041) and higher values for form factor (0.770±0.014). Neurons of atorvastatin-treated birds were significantly more furrowed compared to controls (shape factor: t (12)=2.75, p=0.017, form factor: t (12)=-2.52, p=0.026).

**Figure 5.**
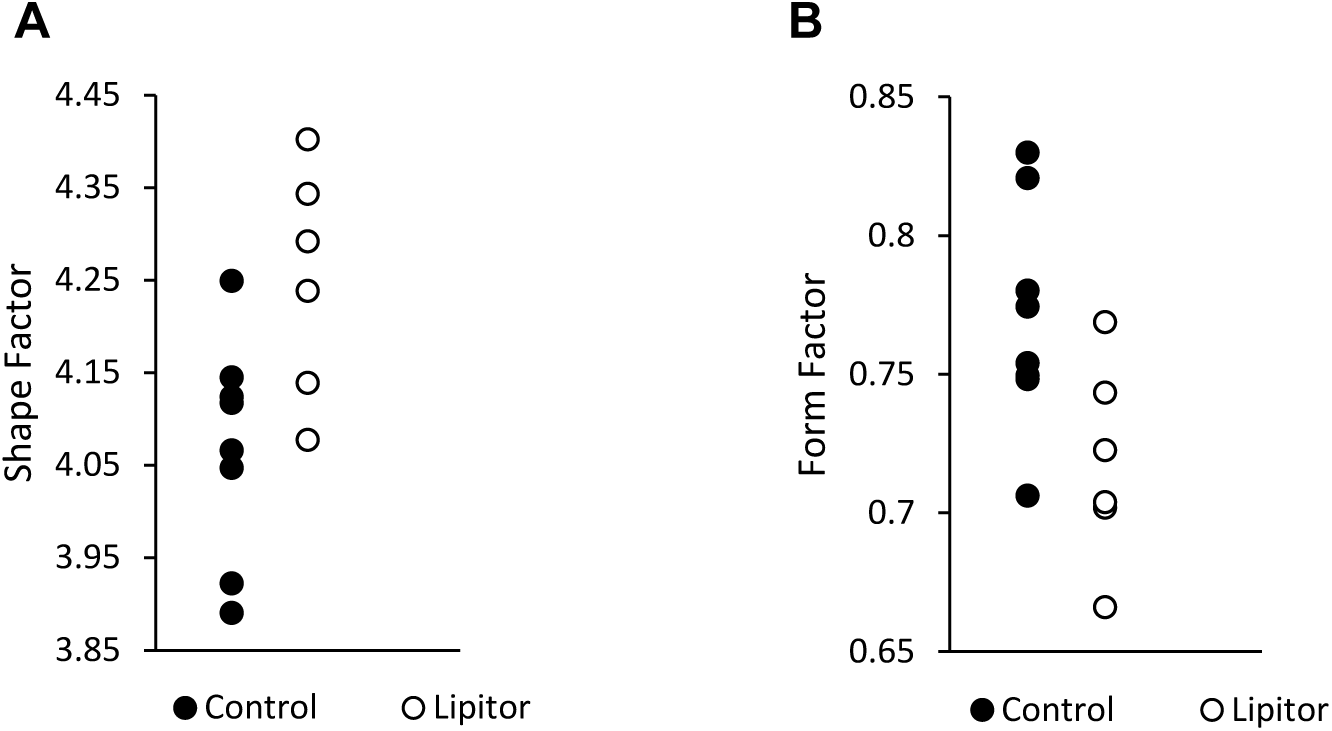
In atorvastatin-treated birds, the contours of ∼30 day old neurons were more furrowed and uneven than in controls. Shape factor and form factor are closely related, both conveying information about an object’s outline. The former is a measure of an object’s area to perimeter ratio, and the latter accounts for both an object’s compactness and also outline complexity. Birds treated with atorvastatin had neuronal soma outlines that were less smooth, and more convoluted as shown by higher shape factor values (A), and low form factor values (B).

### BrdU-/Hu+ cells

We also investigated the morphology of cells expressing the neuronal marker Hu, but not co-labeled with the cell birthdate marker BrdU. For simplicity, we are referring to these as “older” neurons as the majority of these neurons are more likely to be older than 30 days rather than younger, given the limited turnover in HVC and the persistence of long-lived neurons (Walton et al., 2012). Interestingly, unlike the ∼30 day old neurons, atorvastatin treatments did not affect the morphology of older neurons (p > 0.05 for all contour measurements) but had an effect on the neuronal sizes, which were not affected in the ∼30 day old neurons. Older neurons of atorvastatin-treated birds had smaller soma sizes (66.489±3.841) than those of control birds (81.483± 3.621, t (11)=-2.84, p=0.016, Figure 6). The somas of older neurons of atorvastatin-treated birds also had shorter maximum diameters, denoted in Neurolucida as feret max (Figure 7A, 12.678±0.474) than the somas of control birds (13.919±0.227). The somas of older neurons of atorvastatin-treated birds also had shorter minimum diameters, feret min (Figure 7B, 6.994±0.107) than controls (7.774±0.201; Feret Max: t(11)=-2.48, p=0.030, Feret Min: t(11)= - 3.25, p=0.007).

**Figure 6:**
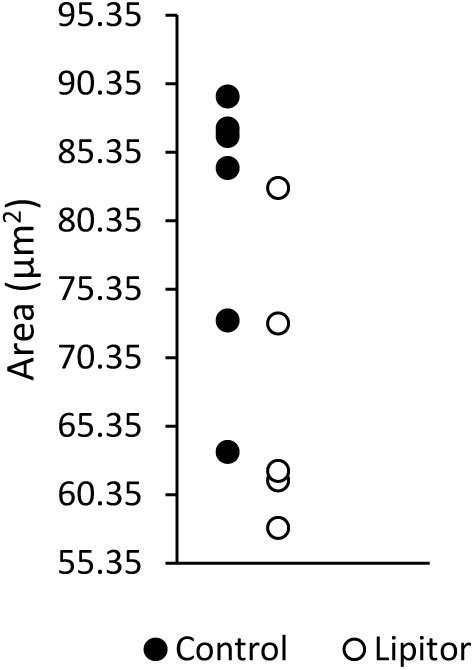
In atorvastatin-treated birds, the soma areas of Hu+ neurons were smaller relative to those of controls.

**Figure 7:**
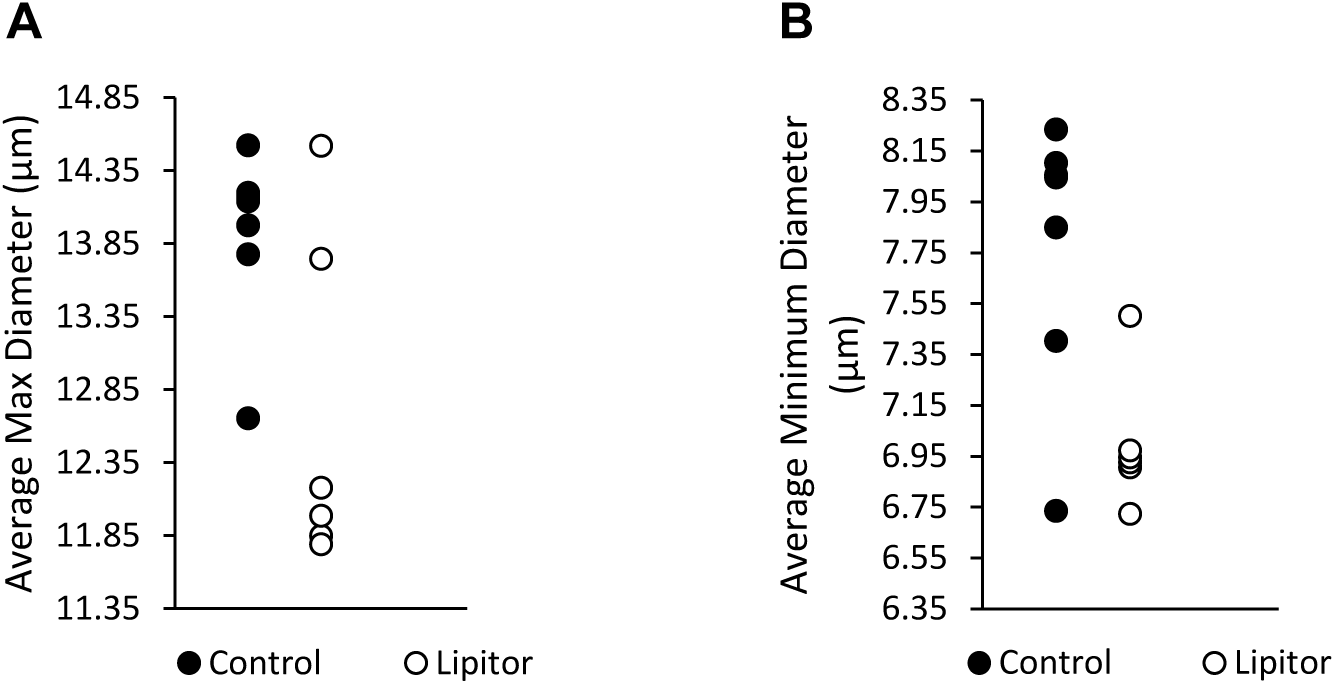
In Atorvastatin-treated birds, the somas of Hu+ cells had shorter maximum and minimum diameters. Neurons not labeled with BrdU, the majority older than ∼30 days, had shorter maximum (A) and minimum diameters (B) in the statin-treated group than in controls.

## Discussion

Animals given a daily oral dose of atorvastatin as juveniles, continuing into adulthood, exhibited numerous morphological differences in neurons of the song motor pathway compared with control birds. New neurons 30-33 days after cell birthdating (BrdU+/Hu+) in treated birds were flatter, had smaller area to maximum diameter ratios, and had furrowed and convoluted outlines relative to those of controls. Neurons of mixed ages (BrdU-/Hu+ cells) did not exhibit any of the morphological differences seen in ∼30 day old neurons. While our sampling of BrdU-/Hu+ neurons likely includes young neurons, we expect that this population primarily consists of older neurons (Walton et al., 2012). We found that BrdU-/Hu+ neurons of birds treated with atorvastatin were smaller than those of controls, as indicated by smaller soma area and shorter soma diameters. New neurons naturally become smaller as they age (Kirn et al., 1991). However, the increased reduction in soma sizes in the statin-treated may be a consequence of decreased cholesterol availability. We found that atorvastatin did not alter the density of new neurons. Taken together, we propose that atorvastatin interfered with the structure of HVC neurons and affected new born neurons by day ∼30 more than existing neurons.

Alternatively, only a subset of HVC neurons are born in adulthood and perhaps this neuronal population is more sensitive to the effects of exposure to atorvastatin, unrelated to being newly formed. New neurons that become part of the motor pathway and project to the robust nucleus of the acropallium (RA) are added throughout the bird’s lifetime, whereas HVC neurons projecting to the anterior forebrain nucleus Area X and interneurons are not. Thus BrdU only labeled HVC-to-RA projection neurons and any feature of these cells, including but not limited to their formation during statin treatment, may have rendered them more susceptible to the effects of statin exposure.

Regardless of the reason for the morphological effects of statins in the BrdU+ neuron group, whether due to neurogenesis or another cell type specific property, we have found that high doses of atorvastatin affect neuronal structure *in vivo*. These results are consistent with findings of structural effects of statin exposure on neurons *in vitro*, although the specific characteristics we describe here have not been reported *in vitro* (Pavlov et al., 1995, Michikawa and Yanagisawa, 1999, Tanaka et al., 2000, Schulz et al., 2004, Marz et al., 2007, Xiang and Reeves, 2009, Mailman et al., 2011).

Statins are competitive inhibitors of HMG CoA reductase, an early enzyme of the cholesterol biosynthesis pathway which catalyzes mevalonate. As a consequence, the production of all intermediate and end products of the mevalonate pathway may be altered by statin treatment (Liao and Laufs, 2005). Three different statins (lovastatin, pravastatin and simvastatin) induced numerous changes in gene expression patterns in the mouse cerebral cortex including genes contributing to cell growth, signaling and trafficking (Johnson-Anuna et al., 2005). Many vitamins, including vitamin A, D, E and K are metabolized by isoprenoid products of the mevalonate pathway, and vitamin E levels have been found to decrease following statin-treatment (Perez-Castrillon et al., 2007, Galli and Iuliano, 2010). Squalene, another mevalonate pathway product, is a precursor for steroid hormones. In the brain, estrogens are important for learning and memory (Remage-Healey et al., 2010, Yoder et al., 2012, Remage-Healey et al., 2013, Remage-Healey, 2014), and also have neuroprotective properties. However, the literature describing the effects of statins on neurohormones is scant and limited to short-term studies, models of disease or in chronically ill patients. Due to the pleiotropic effects of statins, it is difficult to control for many of these factors described above, especially *in vivo*. These findings are a step toward documenting potential neural effects of statin use in the healthy juvenile brain.

## References

Alvarez-Buylla A, Nottebohm F (1988) Migration of young neurons in adult avian brain. Nature 335:353–354.

Banach M, Rizzo M, Nikolic D, Howard G, Howard V, Mikhailidis D (2017) Intensive LDL-cholesterol lowering therapy and neurocognitive function. Pharmacol Ther 170:181–191.

Bunt CW, Hogan AJ (2017) The Effect of Statins on Dementia and Cognitive Decline. Am Fam Physician 95:151–152.

Burns MP, Igbavboa U, Wang L, Wood WG, Duff K (2006) Cholesterol distribution, not total levels, correlate with altered amyloid precursor protein processing in statin-treated mice. Neuromolecular Med 8:319–328.

Chan D, Binks S, Nicholas JM, Frost C, Cardoso MJ, Ourselin S, Wilkie D, Nicholas R, Chataway J (2017) Effect of high-dose simvastatin on cognitive, neuropsychiatric, and health-related quality-of-life measures in secondary progressive multiple sclerosis: secondary analyses from the MS-STAT randomised, placebo-controlled trial. Lancet Neurol 16:591–600.

Chaudhuri A, Chattopadhyay A (2011) Transbilayer organization of membrane cholesterol at low concentrations: Implications in health and disease. Biochim Biophys Acta 1:19–25.

Dietschy JM (2009) Central nervous system: cholesterol turnover, brain development and neurodegeneration. Biol Chem 390:287–293.

Dietschy JM, Turley SD (2001) Cholesterol metabolism in the brain. Curr Opin Lipidol 12:105–112.

Dietschy JM, Turley SD (2004) Thematic review series: brain Lipids. Cholesterol metabolism in the central nervous system during early development and in the mature animal. J Lipid Res 45:1375–1397.

Driver AM, Kratz LE, Kelley RI, Stottmann RW (2016) Altered cholesterol biosynthesis causes precocious neurogenesis in the developing mouse forebrain. Neurobiol Dis 91:69–82.

Eckert GP, Kirsch C, Mueller WE (2001) Differential effects of lovastatin treatment on brain cholesterol levels in normal and apoE-deficient mice. Neuroreport 12:883–887.

Fong CW (2014) Statins in therapy: understanding their hydrophilicity, lipophilicity, binding to 3-hydroxy-3-methylglutaryl-CoA reductase, ability to cross the blood brain barrier and metabolic stability based on electrostatic molecular orbital studies. Eur J Med Chem 85:661–674.

Funfschilling U, Saher G, Xiao L, Mobius W, Nave KA (2007) Survival of adult neurons lacking cholesterol synthesis in vivo. BMC Neurosci 8:1.

Galli F, Iuliano L (2010) Do statins cause myopathy by lowering vitamin E levels? Med Hypotheses 74:707–709.

Gelissen IC, Nguyen HL, Tiao DK, Ayoub R, Aslani P, Moles R (2014) Statin use in Australian children: a retrospective audit of four pediatric hospitals. Paediatr Drugs 16:417–423.

Goluszko P, Nowicki B (2005) Membrane cholesterol: a crucial molecule affecting interactions of microbial pathogens with mammalian cells. Infect Immun 73:7791–7796.

Horwich TB, MacLellan WR, Fonarow GC (2004) Statin therapy is associated with improved survival in ischemic and non-ischemic heart failure. J Am Coll Cardiol 43:642–648.

Johnson-Anuna LN, Eckert GP, Keller JH, Igbavboa U, Franke C, Fechner T, Schubert-Zsilavecz M, Karas M, Muller WE, Wood WG (2005) Chronic administration of statins alters multiple gene expression patterns in mouse cerebral cortex. J Pharmacol Exp Ther 312:786–793.

Kirn JR, Alvarez-Buylla A, Nottebohm F (1991) Production and survival of projection neurons in a forebrain vocal center of adult male canaries. J Neurosci 11:1756–1762.

Kirsch C, Eckert GP, Mueller WE (2003) Statin effects on cholesterol micro-domains in brain plasma membranes. Biochem Pharmacol 65:843–856.

Liao JK, Laufs U (2005) Pleiotropic effects of statins. Annu Rev Pharmacol Toxicol 45:89-118.

Liscum L, Underwood KW (1995) Intracellular cholesterol transport and compartmentation. J Biol Chem 270:15443–15446.

Locatelli S, Lutjohann D, Schmidt HH, Otto C, Beisiegel U, von Bergmann K (2002) Reduction of plasma 24S-hydroxycholesterol (cerebrosterol) levels using high-dosage simvastatin in patients with hypercholesterolemia: evidence that simvastatin affects cholesterol metabolism in the human brain. Arch Neurol 59:213–216.

Mailman T, Hariharan M, Karten B (2011) Inhibition of neuronal cholesterol biosynthesis with lovastatin leads to impaired synaptic vesicle release even in the presence of lipoproteins or geranylgeraniol. J Neurochem 119:1002–1015.

Marz P, Otten U, Miserez AR (2007) Statins induce differentiation and cell death in neurons and astroglia. Glia 55:1–12.

Michikawa M, Yanagisawa K (1999) Inhibition of cholesterol production but not of nonsterol isoprenoid products induces neuronal cell death. J Neurochem 72:2278–2285.

Mouritsen OG, Zuckermann MJ (2004) What's so special about cholesterol? Lipids 39:1101–1113.

Nieweg K, Schaller H, Pfrieger FW (2009) Marked differences in cholesterol synthesis between neurons and glial cells from postnatal rats. J Neurochem 109:125–134.

Orth M, Bellosta S (2012) Cholesterol: its regulation and role in central nervous system disorders. Cholesterol 2012:292598.

Pavlov OV, Bobryshev Yu V, Balabanov Yu V, Ashwell K (1995) An in vitro study of the effects of lovastatin on human fetal brain cells. Neurotoxicol Teratol 17:31–39.

Perez-Castrillon JL, Vega G, Abad L, Sanz A, Chaves J, Hernandez G, Duenas A (2007) Effects of Atorvastatin on vitamin D levels in patients with acute ischemic heart disease. Am J Cardiol 99:903–905.

Pfrieger FW (2003a) Cholesterol homeostasis and function in neurons of the central nervous system. Cell Mol Life Sci 60:1158–1171.

Pfrieger FW (2003b) Role of cholesterol in synapse formation and function. Biochim Biophys Acta 1610:271–280.

Piomelli D, Astarita G, Rapaka R (2007) A neuroscientist's guide to lipidomics. Nat Rev Neurosci 8:743–754.

Pytte CL, George S, Korman S, David E, Bogdan D, Kirn JR (2012) Adult neurogenesis is associated with the maintenance of a stereotyped, learned motor behavior. J Neurosci 32:7052–7057.

Remage-Healey L (2014) Frank Beach Award Winner: Steroids as neuromodulators of brain circuits and behavior. Horm Behav 66:552–560.

Remage-Healey L, Coleman MJ, Oyama RK, Schlinger BA (2010) Brain estrogens rapidly strengthen auditory encoding and guide song preference in a songbird. Proc Natl Acad Sci U S A 107:3852–3857.

Remage-Healey L, Jeon SD, Joshi NR (2013) Recent evidence for rapid synthesis and action of oestrogens during auditory processing in a songbird. J Neuroendocrinol 25:1024–1031.

Saher G, Brugger B, Lappe-Siefke C, Mobius W, Tozawa R, Wehr MC, Wieland F, Ishibashi S, Nave KA (2005) High cholesterol level is essential for myelin membrane growth. Nat Neurosci 8:468–475.

Samaras K, Brodaty H, Sachdev PS (2016) Does statin use cause memory decline in the elderly? Trends Cardiovasc Med 26:550–565.

Schreurs BG (2010) The effects of cholesterol on learning and memory. Neurosci Biobehav Rev 34:1366–1379.

Schulz JG, Bosel J, Stoeckel M, Megow D, Dirnagl U, Endres M (2004) HMG-CoA reductase inhibition causes neurite loss by interfering with geranylgeranylpyrophosphate synthesis. J Neurochem 89:24–32.

Simons K, Ikonen E (2000) How cells handle cholesterol. Science 290:1721–1726.

Suraweera C, de Silva V, Hanwella R (2016) Simvastatin-induced cognitive dysfunction: two case reports. J Med Case Rep 10:016–0877.

Tanaka T, Tatsuno I, Uchida D, Moroo I, Morio H, Nakamura S, Noguchi Y, Yasuda T, Kitagawa M, Saito Y, Hirai A (2000) Geranylgeranyl-pyrophosphate, an isoprenoid of mevalonate cascade, is a critical compound for rat primary cultured cortical neurons to protect the cell death induced by 3-hydroxy-3-methylglutaryl-CoA reductase inhibition. J Neurosci 20:2852–2859.

Theofilopoulos S, Arenas E (2015) Liver X receptors and cholesterol metabolism: role in ventral midbrain development and neurodegeneration. F1000Prime Rep 7:37.

Tsoi SC, Aiya UV, Wasner KD, Phan ML, Pytte CL, Vicario DS (2014) Hemispheric asymmetry in new neurons in adulthood is associated with vocal learning and auditory memory. PLoS One 9:e108929.

Vance JE (2012) Dysregulation of cholesterol balance in the brain: contribution to neurodegenerative diseases. Dis Model Mech 5:746–755.

Vance JE, Hayashi H, Karten B (2005) Cholesterol homeostasis in neurons and glial cells. Semin Cell Dev Biol 16:193–212.

Wagstaff LR, Mitton MW, Arvik BM, Doraiswamy PM (2003) Statin-associated memory loss: analysis of 60 case reports and review of the literature. Pharmacotherapy 23:871–880.

Walton C, Pariser E, Nottebohm F (2012) The zebra finch paradox: song is little changed, but number of neurons doubles. J Neurosci 32:761–774.

Wiegman A, Hutten BA, de Groot E, Rodenburg J, Bakker HD, Buller HR, Sijbrands EJ, Kastelein JJ (2004) Efficacy and safety of statin therapy in children with familial hypercholesterolemia: a randomized controlled trial. JAMA 292:331–337.

Wise LD, Stoffregen DA, Hoe CM, Lankas GR (2011) Juvenile toxicity assessment of open-acid lovastatin in rats. Birth Defects Res B Dev Reprod Toxicol 92:314–322.

Xiang Z, Reeves SA (2009) Simvastatin induces cell death in a mouse cerebellar slice culture (CSC) model of developmental myelination. Exp Neurol 215:41–47.

Yeagle PL (1989) Lipid regulation of cell membrane structure and function. Faseb J 3:1833–1842.

Yoder KM, Lu K, Vicario DS (2012) Blocking estradiol synthesis affects memory for songs in auditory forebrain of male zebra finches. Neuroreport 23:922–926.

